# Different Trajectories of Functional Connectivity Captured with Gamma-Event Coupling and Broadband Measures of EEG in the Rat Fluid Percussion Injury Model

**DOI:** 10.1101/2024.06.02.597056

**Authors:** Rachel Fox, Cesar Santana-Gomez, Mohamad Shamas, Aarja Pavade, Richard Staba, Neil G. Harris

## Abstract

Functional connectivity (FC) after TBI is affected by an altered excitatory-inhibitory balance due to neuronal dysfunction, and the mechanistic changes observed could be reflected differently by contrasting methods. Local gamma event coupling FC (GEC-FC) is believed to represent multiunit fluctuations due to inhibitory dysfunction, and we hypothesized that FC derived from widespread, broadband amplitude signal (BBA-FC) would be different, reflecting broader mechanisms of functional disconnection. We tested this during sleep and active periods defined by high delta and theta EEG activity, respectively, at 1,7 and 28d after rat fluid-percussion-injury (FPI) or sham injury (n=6/group) using 10 indwelling, bilateral cortical and hippocampal electrodes. We also measured seizure and high-frequency oscillatory activity (HFOs) as markers of electrophysiological burden. BBA-FC analysis showed early hyperconnectivity constrained to ipsilateral sensory-cortex-to-CA1-hippocampus that transformed to mainly ipsilateral FC deficits by 28d compared to shams. These changes were conserved over active epochs, except at 28d when there were no differences to shams. In comparison, GEC-FC analysis showed large regions of hyperconnectivity early after injury within similar ipsilateral and intrahemispheric networks. GEC-FC weakened with time, but hyperconnectivity persisted at 28d compared to sham. Edge- and global connectivity measures revealed injury-related differences across time in GEC-FC as compared to BBA-FC, demonstrating greater sensitivity to FC changes post-injury. There was no significant association between sleep fragmentation, HFOs, or seizures with FC changes. The within-animal, spatial-temporal differences in BBA-FC and GEC-FC after injury may represent different mechanisms driving FC changes as a result of primary disconnection and interneuron loss.

**Significance statement:** The present study adds to the understanding of functional connectivity changes in preclinical models of traumatic brain injury. In previously reported literature, there is heterogeneity in the directionality of connectivity changes after injury, resulting from factors such as severity of injury, frequency band studied, and methodology used to calculate FC. This study aims to further clarify differential mechanisms that result in altered network topography after injury, by using Broadband Amplitude-Derived FC and Gamma Event Coupling-Derived FC in EEG. We found post-injury changes that differ in complexity and directionality between measures at and across timepoints. In conjunction with known results and future studies identifying different neural drivers underlying these changes, measures derived from this study could provide useful means from which to minimally-invasively study temporally-evolving pathology after TBI.

## Introduction

Traumatic brain injury (TBI) causes large-scale circuit reorganization with increased local input and disconnect from distant regions^1^. Existing methods to measure functional connectivity (FC) derived from electroencephalographic (EEG) measurements have used phase coupling described by phase coherence, amplitude synchrony described by power of neuronal firing, and cross-frequency coupling between different spectral frequencies^2^. These methods have been applied to EEG and some imaging readouts to generate measures of global and regional connectivity changes after TBI^3–5^. Despite evidence of sensitivity of EEG measures reflecting altered connectivity, few preclinical studies have used EEG after TBI to examine spatial changes in FC^6–7^. Therefore, we designed this study to further clarify neuronal alterations after TBI in the rat fluid percussion model using an EEG network-based approach.

In this study, two methods of FC were used to evaluate cortical network changes: Broadband Amplitude-Derived Functional Connectivity (BBA-FC) at 0-60Hz to reflect broader multi-unit activity within the corresponding region, and Gamma Event Coupling-Derived Functional Connectivity (GEC-FC) to reflect local connectivity measured using the low gamma band (30-55Hz)^2^. GEC-FC reflects gamma activity as a series of gamma events driven by the multi-unit firing of excitatory and inhibitory neurons, calculating the synchronicity between regions as a measure of FC^8^. Physiological alterations occurring after TBI, including altered levels of GABA and interneuron axotomy as a result of metabolic stress, have been hypothesized to interfere with normal gamma oscillations^9–10^. Interneurons are also known to be particularly vulnerable to loss or damage after injury, where the resulting alterations in excitatory-inhibitory balance are potentially implicated in gamma oscillatory changes^10–12^.

TBI has additionally been associated with altered sleep structure, particularly EEG power and timing of stage transitions, as well as increased Sleep Fragmentation^13–15^. Both preclinical and clinical studies have demonstrated correlations between sleep fragmentation and regional connectivity, where disrupted correlations in theta-delta coupling occur after TBI^15–17^. This study aims to further examine whether sleep has a modulatory effect on FC changes post-injury.

Although it is known that TBI results in altered network connectivity, there is heterogeneity in reported local and global changes over time. The method of evaluating connectivity could further reflect different mechanistic changes that occur in response to injury. In this exploratory study we hypothesized that measurement of local GEC-FC and widespread BBA-FC in rats after fluid percussion injury would produce different patterns of connectivity, reflecting broader mechanisms that result in altered network topography related to the primary injury. We also aimed to investigate the relationship between the clinical biomarker of sleep-wake disturbances and FC. These hypotheses were tested during sleep and active periods defined by high delta and theta EEG activity at 1, 7, and 28 days post injury after adult rat, fluid-percussion injury.

## Materials and Methods

### Experimental Protocol

Continuous EEG recordings (24hrs/day) were captured at 1, 7, and 28 days in a Fluid Percussion Injury (FPI) model of TBI in adult rats (n=6/group), for use in calculations of FC. Adult male Sprague-Dawley rats (300-350 g at the time of TBI) obtained from Charles River Sprague Dawley, USA, were housed in pairs under controlled environmental conditions (temperature of 22 ± 1 °C, humidity 30-70%, lights on 06:00-18:00 h) and with pellet food and water provided *ad libitum* (LabDiet, 5001, USA). All experimental procedures involving rats were conducted in accordance with the guidelines and regulations set forth by the University of California Los Angeles Institutional Animal Care and Use Committee.

### Surgical Procedure for TBI Induction and Electrode Implantation

TBI was induced using the lateral fluid percussion animal model^18^. Briefly, rats were anesthetized using 5% isoflurane vaporized in room air, and then maintained at 1.9% throughout the surgery. Following a longitudinal skin incision and clearance of the periosteum, a 5-mm diameter craniectomy was made using a hand-held trephine, centered at −4.5 mm behind bregma and 2.5 mm from the midline over the left cortex, keeping the dura intact. A modified Luer-Lock syringe cap was placed inside the craniectomy and anchored in place with dental acrylate and anchor screw. Injury was induced using a fluid-percussion device equipped with a straight-tip attachment (AmScien Instruments, Model FP 302, Richmond, VA, USA,). The pressure level was adjusted to produce a severe TBI with a target mortality between 20 and 30% within the first 48 h (n=6)^19^. Sham-operated experimental controls (n=6) underwent the same surgical procedures as the TBI rats except for injury induction. The rat was removed from the device and placed on a heating pad immediately after the impact.

A combination of paired intracerebral microwires and epidural screw electrodes were implanted after injury during the same surgery session. Bipolar electrodes with tip separation of 0.5 mm were implanted through a burr hole into the perilesional cortex frontoparietal (AP = −1.7, ML = 4.0 and D = −1.8) and posterior to the craniectomy (AP = −7.6, ML = 4.0 and D = −1.8). An additional bipolar electrode with tip separation of 1.0 mm was implanted in the anterior ipsilateral hippocampus, aiming at the distal CA1 (AP = −3.0, ML = 1.4 and D = −3.6). Four epidural screw electrodes were implanted into ipsilateral fronto-parietal (AP: −1.7; ML: 2.5), contralateral fronto-parietal (AP: 1.7; ML: 2.5), ipsilateral occipital (AP: −7. 6; ML: 2.5) and contralateral occipital (AP: −7.6; ML: 2.5) area^20^. Ground and reference electrodes (stainless steel screws) were inserted in the skull bone overlying the cerebellum (**Fig. 2**).

### EEG Acquisition

Immediately following surgery, rats were placed in a recovery cage with homoeothermic monitoring for recovery. EEG recording began within an hour after completion of the surgery (about 2h following impact) and was performed continuously 24 h/day for the first week after TBI and at 28 days post-injury, with recordings captured at 1, 7, and 28 days post-injury (dpi). EEG was sampled at a minimum of 2 kHz per channel and minimum passband between 0.1 Hz and 1 kHz. EEG was recorded referentially to a screw electrode positioned in the skull overlying the right cerebellar cortex.

### Gamma Event Coupling and Broadband Amplitude FC Analysis

The broader 0-60Hz frequency band was utilized in BBA-FC analysis while the low gamma frequency band 30-55Hz was utilized in GEC-FC analysis, corresponding with existing literature^8,21–22^ (**Fig. 1**). Prior to artifact detection, the EEG data underwent two pre-processing steps to ensure accurate analysis. First, the mean signal across all channels was removed to eliminate any baseline drift or offset, and a notch filter was applied to eliminate powerline noise at 60 Hz. The signal was segmented into shorter windows, typically 0.25s long across all channel to compute, the power in the frequency band of interest (200-600 Hz). This frequency band was chosen due to its relevance in capturing fast artifact activity commonly observed in EEG recordings. Subsequently, the power values were normalized and thresholds are determined based upon the standard deviation of the signal and the percentage of samples above the mean. Artefacts were then identified based on these threshold criteria. Specifically, segments exhibiting power levels above the calculated thresholds were flagged as containing artifacts. Additionally, a zero-crossing criterion was applied to detect rapid changes in the EEG signal, which can be indicative of artifact presence. Segments failing to meet both threshold and zero-crossing criteria were marked as artifacts. To enhance the efficiency of artifact detection, channels were grouped based on their standard deviation. This grouping strategy allows for the identification of artifacts that are present across multiple channels within the same group. Finally, common artifacts detected within the same group were combined and added to the common artefacts in all other groups. This combination method improves the robustness of artifact detection and reduces the risk of false positives or missed detections. The code used for artefact detection is freely available: https://github.com/MohamadShamas/Artefact-Detector.git. For the TBI group, the algorithm removed an average of 17.3 ± 7.9 minutes, corresponding to 39 ± 18.4% of the total raw data file duration. For the Sham group, the artifact removal pipeline reduced an average of 15.6 ± 8.5 minutes, corresponding to 38 ± 20.5% of the total raw data file duration. EEG signal data were then divided into 30s epochs and sleep-stage scored for the 24-hour period at 1, 7, and 28dpi. To maintain consistency between subjects, the first 20 minutes of recordings predominantly defined by theta frequencies for active states or delta frequencies for sleep states at 1, 7, and 28dpi from each of the light and dark periods was used in BBA-FC and GEC-FC calculations and mapped onto a rat brain surface. (**Fig. 2A**). GEC-FC was computed using prior published methods^8,21^ and implemented in Matlab (The Math Works, Inc. MATLAB. Version 2020a).

**Fig. 1.**
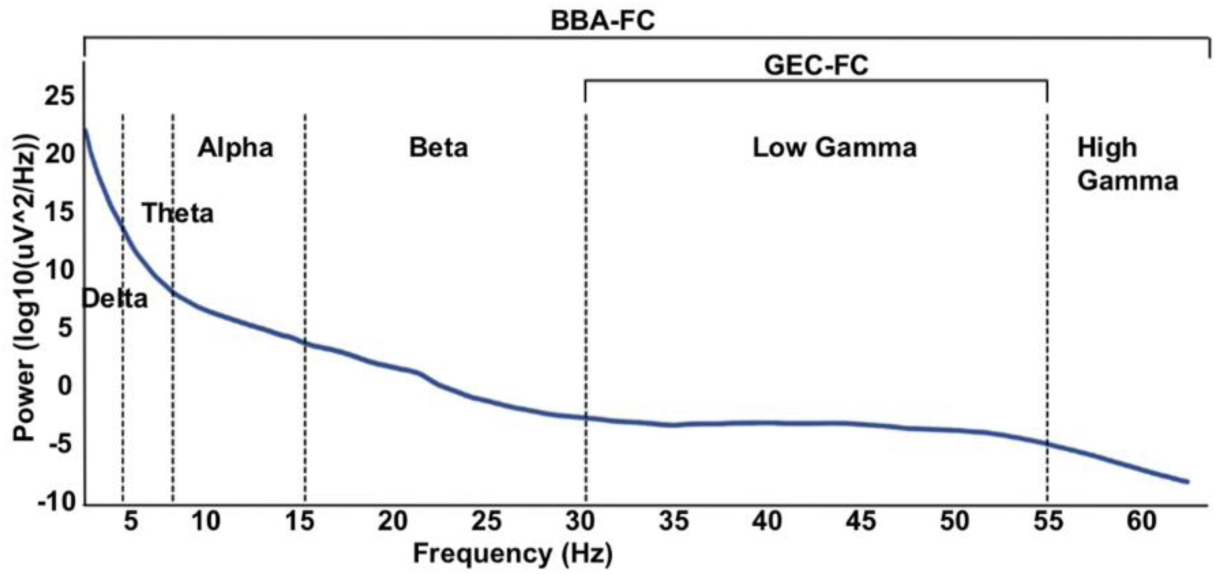
Representative diagram of EEG Power Spectrum with labeled frequency subset used for Broadband Amplitude-Derived Functional Connectivity (BBA-FC, 0-60Hz) and Gamma Event Coupling-Derived Functional Connectivity (GEC-FC, 30-55Hz).

**Fig. 2.**
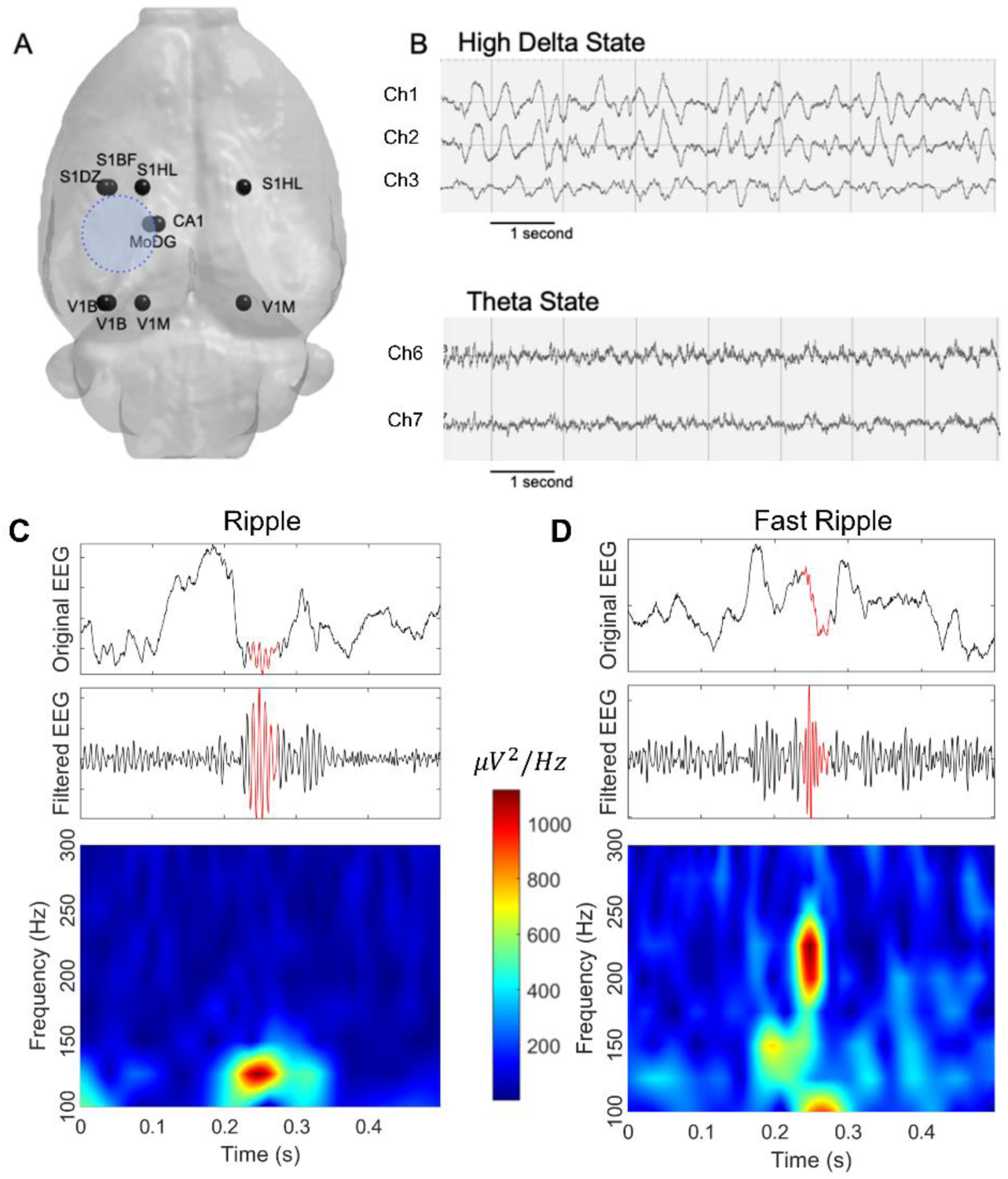
Representative Electrode Placement, EEG Traces, and HFO Analysis: [A] Map of electrode placements (black spheres) and the approximate injury location (blue shaded circle) [B] Representative EEG traces of delta and theta states used in the 24-hour sleep stage scoring and functional connectivity analysis. [C-D] Examples of ripple [C] and Fast Ripple [D] activity, distinguishable by their unique patterns of oscillation, as accentuated by the accompanying heat maps.

Functional connectivity was assessed through the calculation of peri-event histograms, which depicted the rate and timing of events relative to each other. Gamma event coupling is stable across behavioral states, and unlike other phase-amplitude coupling methods, does not have a prior assumption of a predefined model. Each event in the reference channel was analyzed for the occurrence of events in the target channel within a 630-millisecond window. This process was repeated for all detected events in the reference channel, generating a time histogram. To ensure accurate representation, a time resolution of 2 milliseconds was chosen, based on the Nyquist rate principle. The time window for related events was determined as 1 divided by the minimum frequency in the selected frequency band, resulting in a window of less than 34 milliseconds for a minimum frequency of 30 Hz. To generate reliable histograms, a minimum of 30 data points per bin interval was required, necessitating the collection of at least 1020 events within a file of minimum duration of 24 seconds.

Functional connectivity between two channels was inferred when a significant peak was observed in the peri-event histogram. The strength of this peak was quantified using Shannon entropy (S), where lower S values indicated greater certainty and stronger connectivity between the channels involved. A connectivity index (hij) ranging from 0 to 1 was defined to represent the strength of connectivity between two recording channels, with 1 indicating full connectivity and 0 indicating complete disconnection. This index was calculated by subtracting the Shannon entropy of the peri-event histogram between the channels (sij) from its maximum (Smax), then dividing by Smax.

All connectivity indexes were organized into a symmetric connectivity matrix, where each component reflected the connectivity index between different recording sites. This matrix, with dimensions MxM for M recording channels, provided a comprehensive overview of the strength of connectivity between all pairs of recording channels. While GEC-FC was derived from the value of entropy resulting from the peri-event histogram between the peaks of local gamma (30-55 Hz) recorded from each pair of electrodes, BBA-FC was derived from the Pearson correlation coefficient between broadband signals (0.1-60 Hz) recorded from pairs of electrodes.

### Sleep Fragmentation and HFO Analysis

Sleep Fragmentation (SF), which is hypothesized to be a clinical biomarker of recovery after TBI, is measured by the sleep fragmentation index, representing the total number of awakenings and sleep stage shifts divided by total sleep time^23–24^. 24-hour EEG recordings from 1, 7, and 28 dpi were manually scored for wakefulness, high delta, and theta activity using the AASM Sleep-Scoring Guidelines^25^ (**Fig. 2B**). SF was compared across groups and time using a Linear Mixed Model analysis (SPSS Statistical Software, version 29.0). To determine if local network activity had an effect on observed FC changes, 30 minutes/day of artifact-free EEG slow activity during the light period across each of the first 7 days post injury were selected for HFO detection. A computer-based analysis of HFOs was performed on cleaned data using the Matlab-based RippleLab software^26^. The Short-Time Energy (STE) algorithm was used to detect ripples and fast ripples with the following parameters: bandpass filter 80-200Hz (ripples) and 200-500Hz (fast ripples), successive root mean square values greater than 5 standard deviations above the overall root mean square mean within the 3 millisecond window, and minimum duration of 6 milliseconds with more than 8 peaks greater than 2 standard-deviations above the mean value of the band-passed signal^27^. After the automated HFO detection process, all detected events were reviewed manually with ripples or fast ripples centered within a 500 msec duration window with unfiltered EEG signal above the bandpass filtered signal and in adjacent panel, a spectrogram reflecting power of ripples (80-200 Hz) or fast ripples (200-500 Hz) with respect to time. The only events removed were those corresponding with residual artifact (i.e., brief episodes of low-amplitude, high-frequency muscle activity) that were missed during the artifact removal stage. (**Fig. 2C-D**).

### EEG Seizure detection

Continuous EEG recordings from each day were manually reviewed for the presence EEG seizure patterns, defined by Kane et al. for the International Federation of Clinical Neurophysiology as “repetitive epileptiform EEG discharges >2 Hz and/or characteristic pattern with quasi-rhythmic spatiotemporal evolution (i.e. gradual change in frequency, amplitude, morphology and location), lasting at least several seconds (usually >10 sec)”^28^.

Sham-operated rats undergo a craniotomy, a procedure known to potentially induce injury^29,30,31^, and subsequently receive an electrode implantation. Despite the meticulous execution of these surgical procedures, the trigger of acute seizures has been reported in Sham-operated experimental controls equipped with both epidural and intracerebral electrodes, and even in previously Naïve animals with the same electrodes^32^. However, unlike TBI rats in sham conditions, the seizure incidence is markedly low, and seizures typically subside within 72 hours of injury, and almost never develop PTE. The occurrence of post-implantation seizures remains largely unexplored, possibly due to the fact that most studies delay electrographic recordings until 1 week after implantation to minimize the potential effects of surgical stress on EEG.

### Statistics

Group differences in edge connectivity graph measures were calculated using Network Based Statistics (NBS), which is based on the principles of cluster-based thresholding by treating links within networks as connected components^33^. Alterations in functional connectivity were determined using the strength of edge differences between electrodes, as calculated by mean differences of connectivity at each pairwise region of the connectivity matrix. This was evaluated at P<0.01, 2-tailed, using non-parametric permutation testing over 1000 iterations under GraphVar^34^ (version 2.03a). Calculation of persistence of FC differences across time was conducted using linear mixed model analysis of variance (ANOVA) to test for the effect of time, correcting for multiple comparisons, with rat number as a random effect. Preliminary analysis demonstrated no significant difference between FC at delta and theta states, so these periods were combined for all analyses. NBS generally resulted in 1-3 different subnetworks in the different statistical contrasts run. These results were displayed over a brain mesh using the BrainNet Viewer toolbox^35^ (version 1.7). Measures of global connectivity (global efficiency, clustering coefficient, and characteristic path) were derived from the NBS results and compared between TBI and Sham using a 1-way repeated-measures ANOVA with post-hoc testing for group differences at each level of time corrected for multiple comparisons.

## Results

### Differences in Cross-Sectional Functional Connectivity by BBA and GEC Analysis

Cross-sectional analysis of FC metrics at 1, 7, and 28 days post-injury compared to sham revealed only minor significant changes in BBA-derived connectivity. This was composed of ipsilateral sensory-cortex-to-CA1 hippocampus hyperconnectivity at 1 and 7d which transformed to ipsilateral deficits within the ipsilateral visual cortex and bilateral sensory cortex at 28dpi (**Fig. 3A**). There was not a significant effect of time on BBA-FC connections (p>0.05, mixed model repeated measures). In comparison, GEC analysis demonstrated much larger regions of early and persistent ipsilateral hyperconnectivity that weakened over time (p<0.05, mixed model repeated measures ANOVA after correcting for multiple comparisons), but remained significantly hyperconnected within the contralateral sensory-cortex-CA1 and ipsilateral visual cortex compared to shams (**Fig. 3B**). Furthermore, the number of significant edges was greater in GEC-FC compared to BBA-FC at 1 and 7dpi. The average percent difference in FC for significant edges between TBI and Sham was greater in GEC-FC at 1 and 28dpi (100%, 15%, and 153% increase for GEC-FC and 56%, 35% and −4% changes for BBA-FC at 1, 7, 28dpi respectively, two tailed t-test, p<0.05). The difference in directionality of FC changes due to injury could indicate increased sensitivity to underlying pathobiology.

**Fig. 3.**
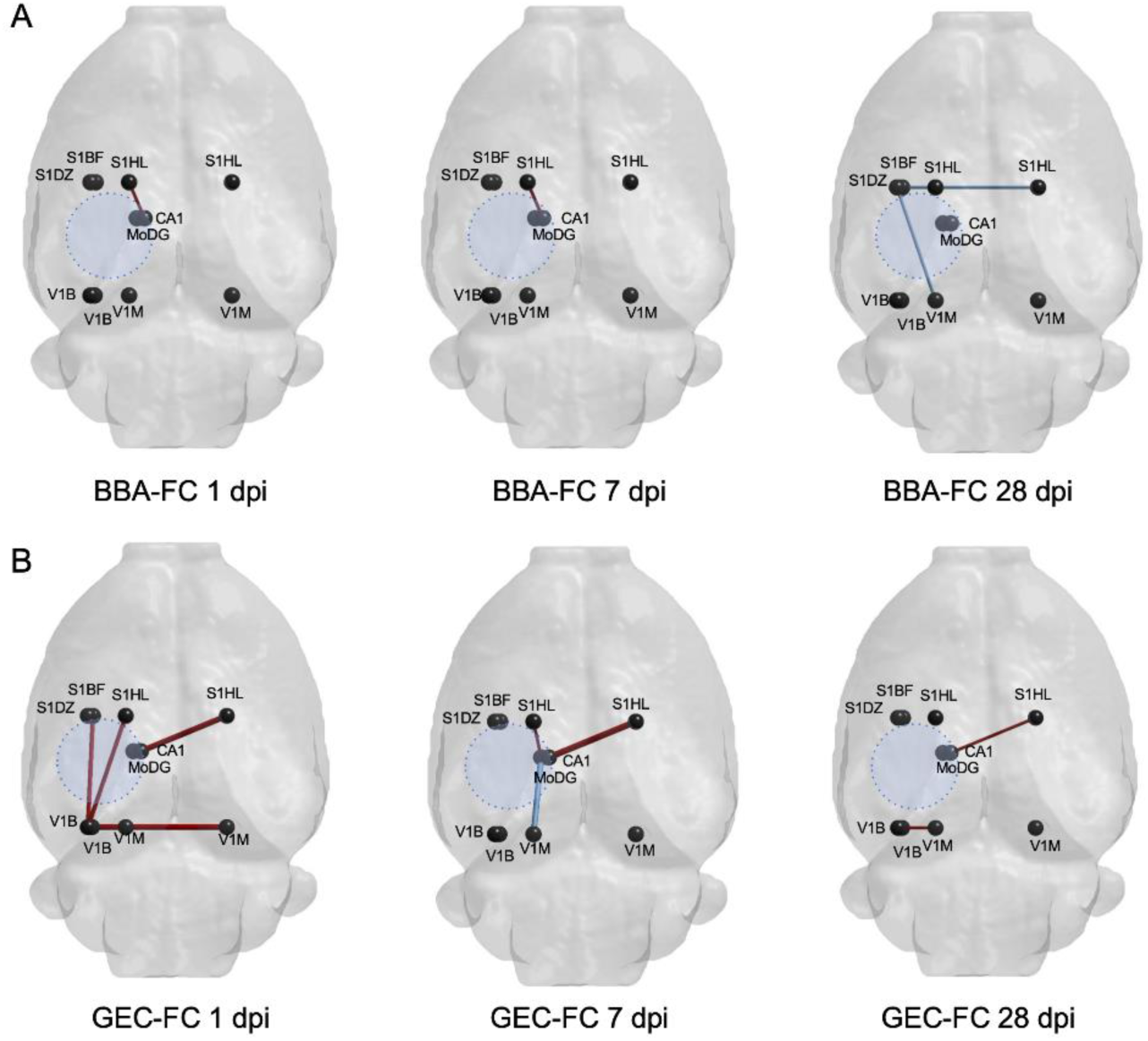
Statistically different edge connectivity between electrodes comparing TBI and sham groups at 1, 7, and 28dpi for: [A] BBA-FC and [B] GEC-FC networks. <u>KEY</u>: Red: FC is TBI>Sham, Blue: FC is TBI<Sham (p<0.01, NBS); Black spheres = electrode positions.

Graph-based network analysis at each timepoint demonstrated a significant effect of day post-injury for the global connectivity metrics characteristic path, clustering path, and global efficiency (1-way repeated measures ANOVA). Post-hoc tests corrected for multiple comparisons revealed significant group level differences only in clustering coefficient measured by GEC-FC at 1dpi (**Fig 4**). Although the magnitude of the BBA-FC global connectivity differences was larger than for GEC-FC, the variance of these group-level differences was greater, possibly contributing to the non-significant differences observed.

**Fig. 4.**
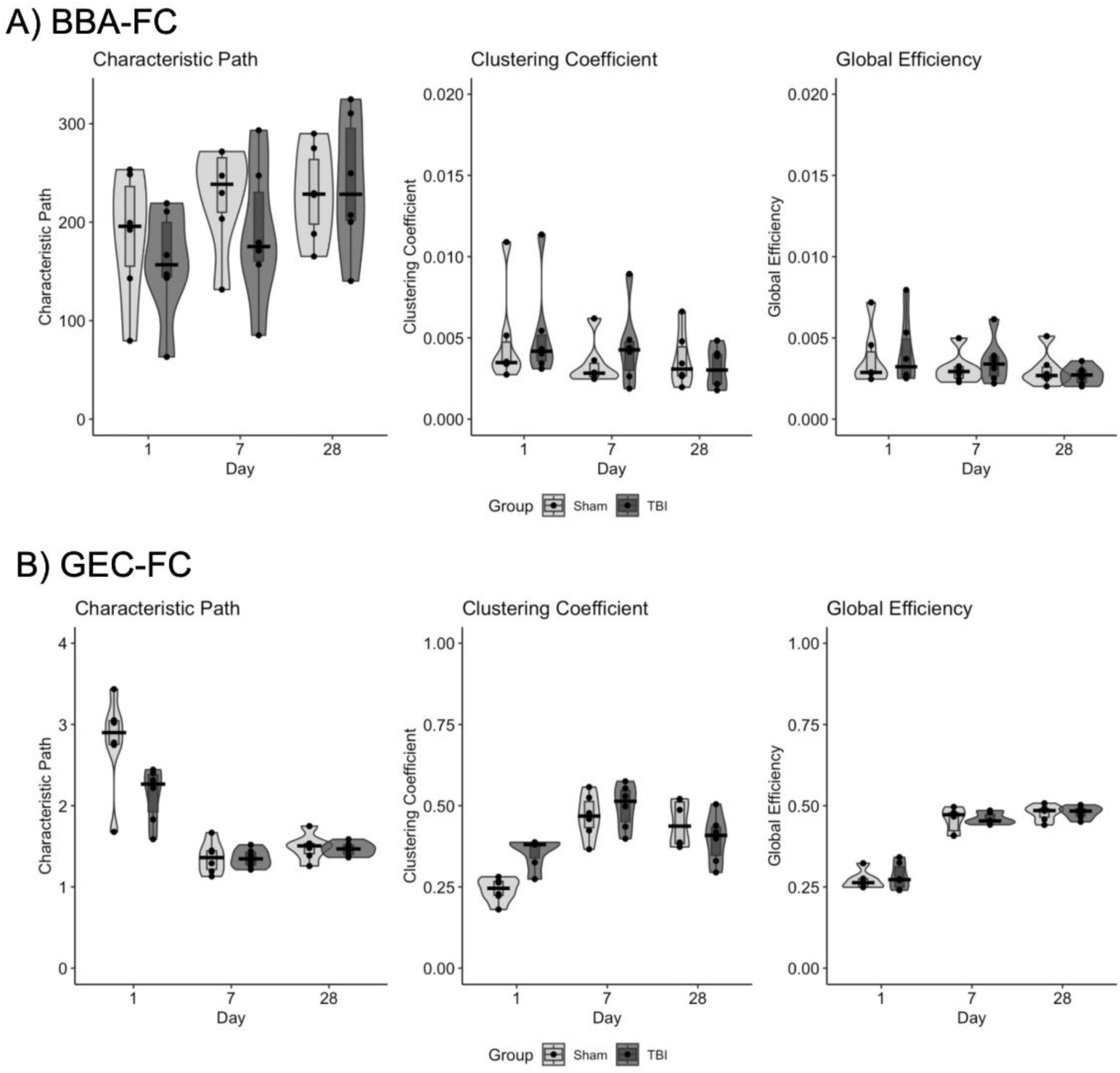
Global Connectivity Measures: Global connectivity measures of characteristic path, clustering coefficient, and global efficiency measured for [A] BBA-FC and [B] GEC-FC analysis. *=P<0.05.

### Temporal Changes in Functional Connectivity by BBA and GEC Analysis

In order to compare the difference in the magnitude of FC change over time between TBI and sham rats, a longitudinal analysis was conducted for each timepoint combination (**Fig. 5A-B**). FC within the Sham group did not differ significantly across time (P>0.05), but there was a significant change of FC in the TBI group across all time-points for both GEC-FC and BBA-FC as compared to Sham (P<0.01). The number of significantly different edges due to injury obtained by GEC-FC analysis was greater than for BBA-FC for all timepoints, both within and between hemispheres, demonstrating a persistence in the extent of network complexity from GEC-FC (**Fig. 5A-B**). The percent change in FC in TBI rats over time using GEC-FC was significantly greater between 1-7dpi and 1-28dpi than BBA-FC analysis, but not between 7-28dpi (317%, 275%, and 511% increase for GEC-FC, versus 208%, −348%, and 87% for BBA-FC, respectively for 1-7, 7-28, and 1-28dpi, two tailed t-test, p<0.05). The temporal decrease in BBA-FC due to injury between days 7-28 corresponded to persistent hypoconnectivity seen at 28dpi obtained by cross-sectional analysis (**Fig. 3A)**.

**Fig 5.**
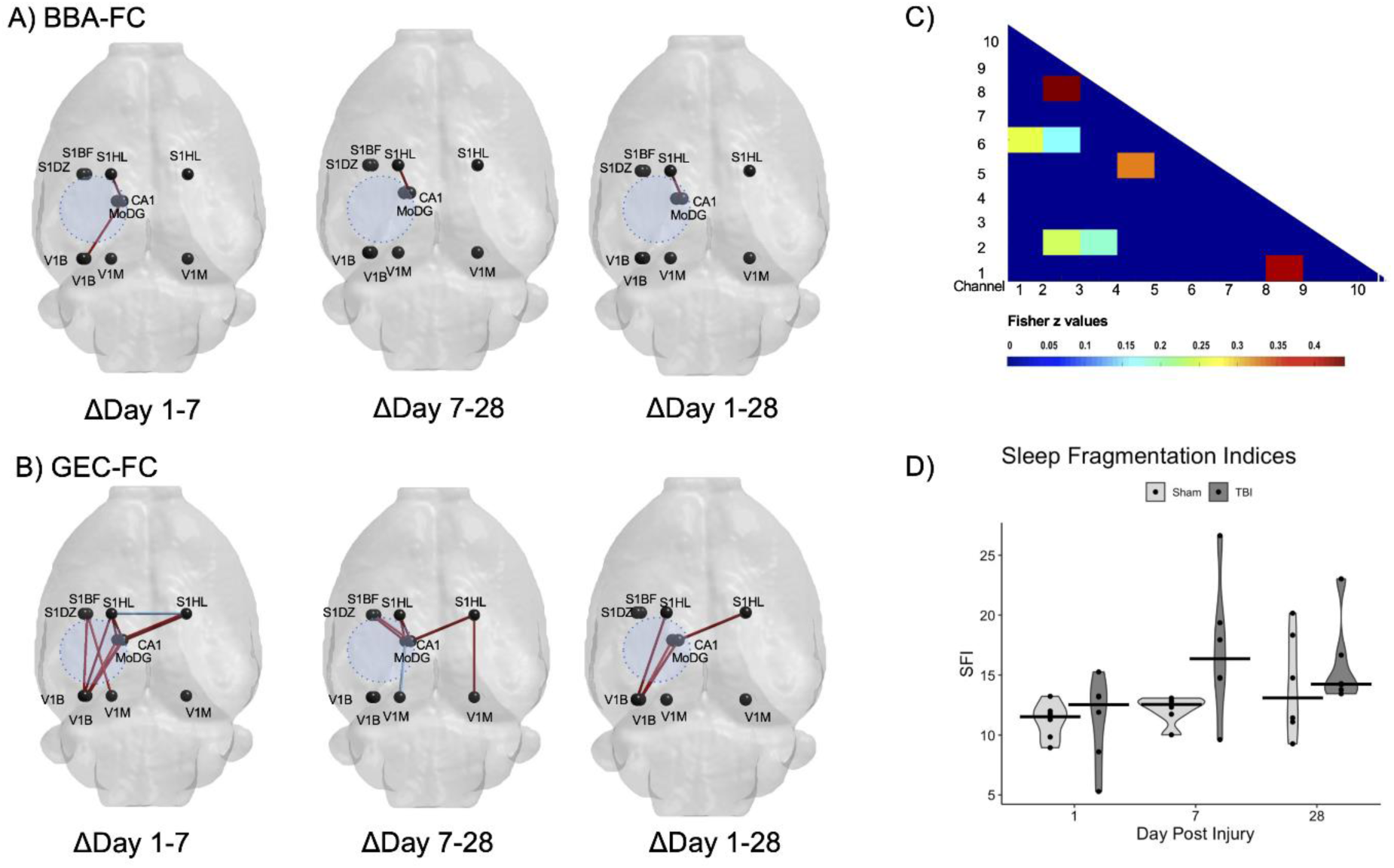
Longitudinal analysis of the strength of FC over time post-injury. The magnitude of [A] BBA-FC and [B] GEC-FC change over time was greater in the TBI group than the Sham group, particularly across hemispheres (p<0.01). [C] Representative matrix of GEC-FC differences, colored boxes indicate significant edge connections between indicated electrodes. [D] Sleep Fragmentation (SF) was significantly increased (p<0.05) in TBI rats at 7dpi and trended towards an increase over time. There was no significant correlation between degree of SF and FC changes. <u>KEY</u>: Red: Increase in FC over time, where TBI >Sham. Blue: Decrease in FC over time, where TBI < Sham (p<0.01, NBS).

Graph-based network analysis of this data revealed a significant injury-related temporal change in GEC-FC-derived global clustering coefficient (two-tailed t-test, p<0.05; **Fig. 4**), but no change in any other measures. There were no significant injury-related temporal changes in any of the network measures derived from the BBA-FC data.

### Sleep Analysis and Electrographic Measures of Local Network Activity

Sleep fragmentation index (SFI) was calculated to test for an association between sleep-wake disturbances and FC. First, we used a linear mixed model analysis with fixed effects and found a significant effect of time and group x time on sleep fragmentation index (p<0.001), but not an effect of group when collapsing across time (**Fig. 5D**). Pairwise comparisons over time revealed significantly higher SFI in TBI compared to sham rats only at 7dpi (p<0.05). We directly tested for a relationship between SFI and FC using SFI as a covariate of interest within the general linear model framework. We found no association between SFI and FC computed with both methods.

HFO ripples, fast ripples, and number of seizures were calculated between 1-7dpi to determine if high frequency activity was present and related to FC changes. Five of the 6 TBI animals developed seizures during the first 7 days after injury, compared with 3 of 6 animals from the Sham group. Although TBI rats had a total of 123 seizures during the acute period as compared to 8 in the sham group, there was no significant difference in seizure count as a result of injury, due to the large variance from one TBI rat with 61 and another with 41 of those 123 seizures. There were also no significant differences in HFO ripples and fast ripples between TBI and sham, suggesting that the FC changes tested for were unconfounded by local network activity or surgery-related effects. Observed FC changes in these data were therefore more likely to be related to the effect of injury alone.

## Discussion

In this study, we found that EEG-derived FC is altered after TBI in the direction of hyperconnectivity at acute timepoints, especially for GEC-derived FC. Significant hyperconnectivity persisted for GEC-FC, while hypoconnectivity was indicated at 28dpi for BBA-FC. Sham FC remained stable over time for both measures adding credence to the injury-related findings.

Altered structural and functional connectivity after TBI are generally accepted as hallmark findings of the spectrum of TBI injuries. Hyperconnectedness has been electrographically associated with a decrease in inhibitory synaptic input, increased spiking activity in the gamma band, and increased local synchrony^21,36–38^. Previous studies have similarly described increased connectivity in the perilesional cortex after injury, namely in regions that were found to undergo structural and circuit reorganization^12,39^. However, other studies have reported heterogeneity in connectivity after TBI, hypothesized to be explained by mechanisms such as variation in injury severity, compensatory reorganization to offset functional loss, and non-uniform expression of hyperconnectivity within subnetworks^39–42^. In this study, GEC-FC captured greater injury-related differences than BBA-FC, as indicated in global connectivity measures across time, a higher number of local subnetwork edges compared to sham, and a smaller variance between subjects. Mechanisms underlying GEC-FC and BBA-FC changes have been hypothesized to be related to axonal injury leading to suppression of inhibition^43^. However, the very different temporal changes in GEC-FC and BBA-FC described herein point to the existence of different underlying neural substrates. Broadband power reflects multi-unit activity from overall circuitry, as measured by the Pearson correlation of EEG power over the 0-60Hz frequency band^44–45^. In contrast, GEC reflects rhythmic synchronization of neural activity and is calculated using temporal relationships between maxima gamma wave activity between regions^2,46^. Decreased BBA-FC-specific sensitivity for detecting injury-related differences could therefore result from measuring the much wider 0-60Hz spectrum compared to the much narrower range for GEC-FC due to frequency-band specific differences^47^.

GEC-FC has been shown to reflect interneuronal activity, where FC changes are related to interneuron-mediated transfer within the cortex^48^. Fast-spiking interneurons contribute to temporal narrow-band gamma oscillations as well as increased gamma oscillation amplitude^43–44^. Parvalbumin positive GABAergic interneurons are particularly essential for generating gamma rhythms and long range connections^49–50^. TBI-induced damage to fast-spiking interneurons results in increases in cortical excitability while promoting compensatory reorganization^11,44^. Inhibitory interneuronal dysfunction can therefore disrupt gamma oscillations as well as the rhythmic inhibition and synchrony necessary for short- and long-range connectivity^51^. Following TBI in rodents, injury-related loss of inhibitory interneurons and inhibitory transmission drives an increased excitatory:inhibitory ratio of activity that ultimately favors hyperexcitatory activity^10,52^. Although we found no effect of acute seizures and HFOs on FC, which would have corresponded to this expected increased excitatory:inhibitory ratio, it is possible that GEC-FC driven interneuronal changes could be different from those that support seizures and HFOs. GEC-FC could be therefore used as a method to discern cellular causes of network reorganization that is more specific to interneuronal changes. Although interneurons are a suggested contributor to FC changes after TBI, future work is necessary to determine whether a causal association is present.

The small sample size used in this study makes this work exploratory only. The statistical permutation method used enables only a determination of subnetwork-level alterations and not electrode-to-electrode specific changes. Although sleep fragmentation was not revealed as a significant modulator of FC, there were significant variations in SF scores between individual subjects, so it is possible that all effects were not captured due to this limited group size.

In summary, BBA-FC and GEC-FC reveal initial interhemispheric hyperconnectivity after TBI, but display different trajectories of FC abnormalities toward more chronic post-injury timepoints. Evidence points to different neural drivers underlying these changes, suggesting that these measures may provide a useful means by which to minimally-invasively study temporally-evolving pathology after TBI.

## Acknowledgments

A conference abstract (#DB09) on this work has previously been published: https://www.liebertpub.com/doi/10.1089/neu.2023.29130.abstracts

## Authorship contribution statement

**R. Fox:** Formal Analysis, Visualization, Writing. **C. Santana-Gomez:** Data curation, Methodology, **M. Shamas:** Data curation, Software, **A. Pavade:** Data curation, **R. Staba:** Project administration, writing, funding **N.G. Harris:** Conceptualization, Writing, Project administration, funding.

## References

1. Nakamura T, Hillary FG, Biswal BB. Resting Network Plasticity Following Brain Injury. PLOS ONE 2009;4(12):e8220; doi: 10.1371/journal.pone.0008220.

2. Buzsáki G, Wang X-J. Mechanisms of Gamma Oscillations. Annu Rev Neurosci 2012;35:203–225; doi: 10.1146/annurev-neuro-062111-150444.

3. Meningher I, Bernstein-Eliav M, Rubovitch V, et al. Alterations in Network Connectivity after Traumatic Brain Injury in Mice. Journal of Neurotrauma 2020;37(20):2169–2179; doi: 10.1089/neu.2020.7063.

4. Dunkley BT, Da Costa L, Bethune A, et al. Low-frequency connectivity is associated with mild traumatic brain injury. NeuroImage: Clinical 2015;7:611–621; doi: 10.1016/j.nicl.2015.02.020.

5. Cramer SW, Haley SP, Popa LS, et al. Wide-field calcium imaging reveals widespread changes in cortical functional connectivity following mild traumatic brain injury in the mouse. Neurobiology of Disease 2023;176:105943; doi: 10.1016/j.nbd.2022.105943.

6. Paterno R, Metheny H, Xiong G, et al. Mild Traumatic Brain Injury Decreases Broadband Power in Area CA1. Journal of Neurotrauma 2016;33(17):1645–1649; doi: 10.1089/neu.2015.4107.

7. Johnstone VPA, Shultz SR, Yan EB, et al. The Acute Phase of Mild Traumatic Brain Injury Is Characterized by a Distance-Dependent Neuronal Hypoactivity. Journal of Neurotrauma 2014;31(22):1881–1895; doi: 10.1089/neu.2014.3343.

8. Bragin A, Almajano J, Kheiri F, et al. Functional Connectivity in the Brain Estimated by Analysis of Gamma Events. PLOS ONE 2014;9(1):e85900; doi: 10.1371/journal.pone.0085900.

9. Raible DJ, Frey LC, Cruz Del Angel Y, et al. GABAA Receptor Regulation after Experimental Traumatic Brain Injury. Journal of Neurotrauma 2012;29(16):2548–2554; doi: 10.1089/neu.2012.2483.

10. Vascak M, Jin X, Jacobs KM, et al. Mild Traumatic Brain Injury Induces Structural and Functional Disconnection of Local Neocortical Inhibitory Networks via Parvalbumin Interneuron Diffuse Axonal Injury. Cereb Cortex 2018;28(5):1625–1644; doi: 10.1093/cercor/bhx058.

11. Harris AC, Jin X-T, Greer JE, et al. Somatostatin interneurons exhibit enhanced functional output and resilience to axotomy after mild traumatic brain injury. Neurobiol Dis 2022;171:105801; doi: 10.1016/j.nbd.2022.105801.

12. Frankowski JC, Tierno A, Pavani S, et al. Brain-wide reconstruction of inhibitory circuits after traumatic brain injury. Nat Commun 2022;13(1):3417; doi: 10.1038/s41467-022-31072-2.

13. Andrade P, Lara-Valderrábano L, Manninen E, et al. Seizure Susceptibility and Sleep Disturbance as Biomarkers of Epileptogenesis after Experimental TBI. Biomedicines 2022;10(5):1138; doi: 10.3390/biomedicines10051138.

14. Konduru SS, Wallace EP, Pfammatter JA, et al. Sleep-wake characteristics in a mouse model of severe traumatic brain injury: Relation to posttraumatic epilepsy. Epilepsia Open 2021;6(1):181–194; doi: 10.1002/epi4.12462.

15. Iyer KK, Zalesky A, Barlow KM, et al. Default mode network anatomy and function is linked to pediatric concussion recovery. Ann Clin Transl Neurol 2019;6(12):2544–2554; doi: 10.1002/acn3.50951.

16. Lombardi F, Gómez-Extremera M, Bernaola-Galván P, et al. Critical Dynamics and Coupling in Bursts of Cortical Rhythms Indicate Non-Homeostatic Mechanism for Sleep-Stage Transitions and Dual Role of VLPO Neurons in Both Sleep and Wake. J Neurosci 2020;40(1):171–190; doi: 10.1523/JNEUROSCI.1278-19.2019.

17. Lu J, Greco MA, Shiromani P, et al. Effect of Lesions of the Ventrolateral Preoptic Nucleus on NREM and REM Sleep. J Neurosci 2000;20(10):3830–3842; doi: 10.1523/JNEUROSCI.20-10-03830.2000.

18. McIntosh TK, Vink R, Noble L, et al. Traumatic brain injury in the rat: Characterization of a lateral fluid-percussion model. Neuroscience 1989;28(1):233–244; doi: 10.1016/0306-4522(89)90247-9.

19. Pitkänen A, Mcintosh TK. Animal Models of Post-Traumatic Epilepsy. Journal of Neurotrauma 2006;23(2):241–261; doi: 10.1089/neu.2006.23.241.

20. Paxinos, G. and Watson, C. (2007) The Rat Brain in Stereotaxic Coordinates. 6th Edition, Academic Press, San Diego.

21. Shamas M, Yeh HJ, Fried I, et al. Interictal Gamma Event Connectivity Differentiates the Seizure Network and Outcome in Patients after Temporal Lobe Epilepsy Surgery. eNeuro 2022;9(6); doi: 10.1523/ENEURO.0141-22.2022.

22. Slewa-Younan S, Green AM, Baguley IJ, et al. Is ‘gamma’ (40 Hz) synchronous activity disturbed in patients with traumatic brain injury? Clinical Neurophysiology 2002;113(10):1640–1646; doi: 10.1016/S1388-2457(02)00239-0.

23. Fleming MK, Smejka T, Henderson Slater D, et al. Sleep Disruption After Brain Injury Is Associated With Worse Motor Outcomes and Slower Functional Recovery. Neurorehabil Neural Repair 2020;34(7):661–671; doi: 10.1177/1545968320929669.

24. Haba-Rubio J, Ibanez V, Sforza E. An alternative measure of sleep fragmentation in clinical practice: the sleep fragmentation index. Sleep Medicine 2004;5(6):577–581; doi: 10.1016/j.sleep.2004.06.007.

25. Iber C, Ancoli-Israel S, Chesson A, and Quran SF for the American Academy of Sleep Medicine. The AASm Manual for the Scoring of Sleep and Associated Events: Rules, Terminology and Technical Specifications, 1st ed.: Westchester, Illinois: American Academy of Sleep Medicine, 2007.

26. Navarrete M, Alvarado-Rojas C, Quyen MLV, et al. RIPPLELAB: A Comprehensive Application for the Detection, Analysis and Classification of High Frequency Oscillations in Electroencephalographic Signals. PLOS ONE 2016;11(6):e0158276; doi: 10.1371/journal.pone.0158276.

27. Staba RJ, Wilson CL, Bragin A, et al. Quantitative Analysis of High-Frequency Oscillations (80–500 Hz) Recorded in Human Epileptic Hippocampus and Entorhinal Cortex. Journal of Neurophysiology 2002;88(4):1743–1752; doi: 10.1152/jn.2002.88.4.1743.

28. Kane N, Acharya J, Benickzy S, et al. A revised glossary of terms most commonly used by clinical electroencephalographers and updated proposal for the report format of the EEG findings. Revision 2017. Clin Neurophysiol Pract 2017;2:170–185; doi: 10.1016/j.cnp.2017.07.002.

29. Cole JT, Yarnell A, Kean WS, et al. Craniotomy: True Sham for Traumatic Brain Injury, or a Sham of a Sham? J Neurotrauma 2011;28(3):359–369; doi: 10.1089/neu.2010.1427.

30. Wu JC-C, Chen K-Y, Yo Y-W, et al. Different sham procedures for rats in traumatic brain injury experiments induce corresponding increases in levels of trauma markers. Journal of Surgical Research 2013;179(1):138–144; doi: 10.1016/j.jss.2012.09.013.

31. Shoffstall AJ, Paiz J, Miller D, et al. Potential for Thermal Damage to the Blood-Brain Barrier during Craniotomy Procedure: Implications for Intracortical Recording Microelectrodes. J Neural Eng 2018;15(3):034001; doi: 10.1088/1741-2552/aa9f32.

32. Andrade P, Banuelos-Cabrera I, Lapinlampi N, et al. Acute Non-Convulsive Status Epilepticus after Experimental Traumatic Brain Injury in Rats. J Neurotrauma 2019;36(11):1890–1907; doi: 10.1089/neu.2018.6107.

33. Zalesky A, Fornito A, Bullmore ET. Network-based statistic: Identifying differences in brain networks. NeuroImage 2010;53(4):1197–1207; doi: 10.1016/j.neuroimage.2010.06.041.

34. Kruschwitz JD, List D, Waller L, et al. GraphVar: A user-friendly toolbox for comprehensive graph analyses of functional brain connectivity. Journal of Neuroscience Methods 2015;245:107–115; doi: 10.1016/j.jneumeth.2015.02.021.

35. Xia M, Wang J, He Y. BrainNet Viewer: A Network Visualization Tool for Human Brain Connectomics. PLoS One 2013;8(7):e68910; doi: 10.1371/journal.pone.0068910.

36. Nichols J, Perez R, Wu C, et al. Traumatic Brain Injury Induces Rapid Enhancement of Cortical Excitability in Juvenile Rats. CNS Neurosci Ther 2014;21(2):193–203; doi: 10.1111/cns.12351.

37. Uhlhaas PJ, Singer W. Neural Synchrony in Brain Disorders: Relevance for Cognitive Dysfunctions and Pathophysiology. Neuron 2006;52(1):155–168; doi: 10.1016/j.neuron.2006.09.020.

38. Huttunen JK, Airaksinen AM, Barba C, et al. Detection of Hyperexcitability by Functional Magnetic Resonance Imaging after Experimental Traumatic Brain Injury. Journal of Neurotrauma 2018;35(22):2708–2717; doi: 10.1089/neu.2017.5308.

39. Harris NG, Verley DR, Gutman BA, et al. Disconnection and hyper-connectivity underlie reorganization after TBI: A rodent functional connectomic analysis. Experimental Neurology 2016;277:124–138; doi: 10.1016/j.expneurol.2015.12.020.

40. Hillary FG, Rajtmajer SM, Roman CA, et al. The Rich Get Richer: Brain Injury Elicits Hyperconnectivity in Core Subnetworks. PLOS ONE 2014;9(8):e104021; doi: 10.1371/journal.pone.0104021.

41. Huang M-X, Harrington DL, Robb Swan A, et al. Resting-State Magnetoencephalography Reveals Different Patterns of Aberrant Functional Connectivity in Combat-Related Mild Traumatic Brain Injury. Journal of Neurotrauma 2017;34(7):1412–1426; doi: 10.1089/neu.2016.4581.

42. Mishra AM, Bai X, Sanganahalli BG, et al. Decreased Resting Functional Connectivity after Traumatic Brain Injury in the Rat. PLOS ONE 2014;9(4):e95280; doi: 10.1371/journal.pone.0095280.

43. Guyon N, Zacharias LR, Fermino de Oliveira E, et al. Network Asynchrony Underlying Increased Broadband Gamma Power. J Neurosci 2021;41(13):2944–2963; doi: 10.1523/JNEUROSCI.2250-20.2021.

44. Cardin JA, Carlén M, Meletis K, et al. Driving fast-spiking cells induces gamma rhythm and controls sensory responses. Nature 2009;459(7247):663–667; doi: 10.1038/nature08002.

45. Manning JR, Jacobs J, Fried I, et al. Broadband Shifts in Local Field Potential Power Spectra Are Correlated with Single-Neuron Spiking in Humans. J Neurosci 2009;29(43):13613–13620; doi: 10.1523/JNEUROSCI.2041-09.2009.

46. Kheiri F, Bragin A, Jr JE. Functional connectivity between brain areas estimated by analysis of gamma waves. Journal of Neuroscience Methods 2013;214(2):184–191; doi: 10.1016/j.jneumeth.2013.01.007.

47. Lewine JD, Plis S, Ulloa A, et al. Quantitative EEG Biomarkers for Mild Traumatic Brain Injury. Journal of Clinical Neurophysiology 2019;36(4):298; doi: 10.1097/WNP.0000000000000588.

48. Sohal VS, Zhang F, Yizhar O, et al. Parvalbumin neurons and gamma rhythms enhance cortical circuit performance. Nature 2009;459(7247):698–702; doi: 10.1038/nature07991.

49. Veit J, Hakim R, Jadi MP, et al. Cortical gamma band synchronization through somatostatin interneurons. Nat Neurosci 2017;20(7):951–959; doi: 10.1038/nn.4562.

50. Wang X-J, Buzsáki G. Gamma Oscillation by Synaptic Inhibition in a Hippocampal Interneuronal Network Model. J Neurosci 1996;16(20):6402–6413; doi: 10.1523/JNEUROSCI.16-20-06402.1996.

51. Hunt RFI, Boychuk JA, Smith BN. Neural circuit mechanisms of post–traumatic epilepsy. Front Cell Neurosci 2013;7; doi: 10.3389/fncel.2013.00089.

52. Gu F, Parada I, Shen F, et al. Structural alterations in fast-spiking GABAergic interneurons in a model of posttraumatic neocortical epileptogenesis. Neurobiology of Disease 2017;108:100–114; doi: 10.1016/j.nbd.2017.08.008.

